# Shank promotes action potential repolarization by recruiting BK channels to calcium nanodomains

**DOI:** 10.1101/2021.11.05.467415

**Authors:** Luna Gao, Jian Zhao, Evan Ardiel, Qi Hall, Stephen Nurrish, Joshua M. Kaplan

## Abstract

Mutations altering the scaffolding protein Shank are linked to several psychiatric disorders, and to synaptic and behavioral defects in mice. Among its many binding partners, Shank directly binds CaV1 voltage activated calcium channels. Here we show that the *C. elegans* SHN-1/Shank promotes CaV1 coupling to calcium activated potassium channels. Mutations inactivating SHN-1, and those preventing SHN-1 binding to EGL-19/CaV1 all increase action potential durations in body muscles. Action potential repolarization is mediated by two classes of potassium channels: SHK-1/KCNA and SLO-1 and SLO-2 BK channels. BK channels are calcium-dependent, and their activation requires tight coupling to EGL-19/CaV1 channels. SHN-1’s effects on AP duration are mediated by changes in BK channels. In *shn-1* mutants, SLO-2 currents and channel clustering are significantly decreased in both body muscles and neurons. Finally, increased and decreased *shn-1* gene copy number produce similar changes in AP width and SLO-2 current. Collectively, these results suggest that an important function of Shank is to promote nanodomain coupling of BK with CaV1.

## Introduction

Shank is a synaptic scaffolding protein (containing SH3, PDZ, proline-rich and SAM domains) (Grabrucker et al., 2011). Mammals have three Shank genes, each encoding multiple isoforms (Jiang and Ehlers, 2013). Several mouse Shank knockouts have been described but these mutants exhibit inconsistent (often contradictory) synaptic and behavioral defects (Jiang and Ehlers, 2013), most likely resulting from differences in the Shank isoforms impacted by each mutation. The biochemical mechanism by which Shank mutations alter synaptic function and behavior has not been determined.

In humans, Shank mutations and CNVs are linked to Autism Spectrum Disorders (ASD), schizophrenia, and mania (Durand et al., 2007; Peca et al., 2011). Haploinsufficiency for 22q13 (which spans the Shank3 locus) occurs in Phelan-McDermid syndrome (PMS), a syndromic form of ASD (Phelan and McDermid, 2012). PMS patients exhibit autistic behaviors accompanied by hypotonia, delayed speech, and intellectual disability (ID) (Bonaglia et al., 2011). Heterozygous inactivating Shank3 mutations are found in sporadic ASD and schizophrenia (Durand et al., 2007; Peca et al., 2011). A parallel set of genetic studies suggest that increased Shank3 function also contributes to psychiatric diseases. 22q13 duplications spanning Shank3 are found in ASD, schizophrenia, ADHD, and bipolar disorder (Durand et al., 2007; Failla et al., 2007; Han et al., 2013). A transgenic mouse that selectively over-expresses Shank3 exhibits hyperactive behavior and susceptibility to seizures (Han et al., 2013). Taken together, these studies suggest that too little or too much Shank3 can contribute to the pathophysiology underlying these psychiatric disorders.

Given its link to psychiatric disorders, there is great interest in determining how Shank regulates circuit development and function. Shank is highly enriched in the post-synaptic densities of excitatory synapses; consequently, most studies have focused on the idea that Shank proteins regulate some aspect of synapse formation or function. Through its various domains, Shank proteins bind many proteins (Lee et al., 2011; Sakai et al., 2011), thereby potentially altering diverse cellular functions. Shank proteins have been implicated in activity induced gene transcription (Perfitt et al., 2020; Pym et al., 2017), synaptic transmission (Zhou et al., 2016), synapse maturation (Harris et al., 2016), synaptic homeostasis (Tatavarty et al., 2020), cytoskeletal remodeling (Lilja et al., 2017), and sleep (Ingiosi et al., 2019). Each of these defects could contribute to the neurodevelopmental and cognitive deficits observed in ASD and schizophrenia.

Several recent studies suggest that an important function of Shank is to regulate the subcellular localization of ion channels. Shank mutations decrease the synaptic localization of NMDA and AMPA type glutamate receptors (Peca et al., 2011; Won et al., 2012). Other studies show that Shank proteins promote delivery of several ion channels to the plasma membrane, including HCN channels (Yi et al., 2016; Zhu et al., 2018), TRPV channels (Han et al., 2016), and CaV1 voltage activated calcium channels (Pym et al., 2017; Wang et al., 2017). Of these potential binding partners, we focus on CaV1 because human CACNA1C (which encodes a CaV1 *α*-subunit) is mutated in Timothy Syndrome (TS), a rare monogenic form of ASD (Splawski et al., 2005; Splawski et al., 2004), and polymorphisms linked to CACNA1C are associated with multiple psychiatric disorders (Cross-Disorder Group of the Psychiatric Genomics, 2013). For this reason, we asked how Shank regulates the coupling of CaV1 channels to their downstream effectors.

*C. elegans* has a single Shank gene, *shn-1*. The SHN-1 protein lacks an SH3 domain but has all other domains found in mammalian Shank proteins. Mammalian Shank proteins directly bind CaV1 channels through both the SH3 and PDZ domains (Zhang et al., 2005). We previously showed that the SHN-1 PDZ domain directly binds to a carboxy-terminal ligand in EGL-19/CaV1 (Pym et al., 2017). CaV1 channels are tightly coupled to multiple downstream calcium activated pathways. *C. elegans* and mouse Shank proteins have been shown to promote CaV1 mediated activation of the transcription factor CREB (Perfitt et al., 2020; Pym et al., 2017).

Here we test the idea that SHN-1 regulates CaV1 coupling to a second effector, calcium activated potassium currents (which are mediated by BK channels). BK channels are activated by both membrane depolarization and by cytoplasmic calcium. At resting cytoplasmic calcium levels (∼100 nM), BK channels have extremely low open probability. Following depolarization, cytoplasmic calcium rises thereby activating BK channels. BK channels bind calcium with a K_d_ ranging from 1-10 μM (Contreras et al., 2013); consequently, efficient BK channel activation requires tight spatial coupling to voltage activated calcium (CaV) channels. BK channels associate with all classes of CaV channels (Berkefeld et al., 2006). The co-clustering of BK and CaV channels allows rapid activation of hyperpolarizing potassium currents following depolarization. BK channels decrease action potential (AP) durations, promote rapid after hyperpolarizations, decrease the duration of calcium entry, and limit secretion of neurotransmitters and hormones in neurons and muscles (Adams et al., 1982; Edgerton and Reinhart, 2003; Petersen and Maruyama, 1984; Storm, 1987). Thus, BK channels have profound effects on circuit activity.

*C. elegans* has two BK channel subunits (SLO-1 and SLO-2), both of which form calcium and voltage dependent potassium channels when heterologously expressed (Wang et al., 2001; Yuan et al., 2000). SLO-2 channels are also activated by cytoplasmic chloride (Yuan et al., 2000). As in mammals, neuronal SLO-1 and -2 channels inhibit neurotransmitter release (Liu et al., 2014; Liu et al., 2007; Sancar et al., 2011), presumably via their coupling to UNC-2/CaV2 and EGL-19/CaV1. In body muscles, SLO-1 channels are co-localized with EGL-19/CaV1 channels and regulate muscle excitability and behavior (Kim et al., 2009). Here we show that SHN-1 promotes BK coupling to EGL-19/CaV1 channels, thereby decreasing AP duration.

## Results

### SHN-1 acts in muscles to regulate action potential duration

EGL-19/CaV1 channels mediate the primary depolarizing current during body muscle APs (Jospin et al., 2002; Liu et al., 2011). Because SHN-1 directly binds EGL-19 (Pym et al., 2017), we asked if SHN-1 regulates muscle AP firing patterns. In WT animals, body muscles exhibit a pattern of spontaneous AP bursts (∼10 APs/burst; burst frequency 0.1 Hz) (Fig. 1A). Within a burst, APs became progressively wider (Fig. 1B). A similar pattern of progressive AP broadening during burst firing has been reported for many neurons (Geiger and Jonas, 2000; Jackson et al., 1991). Outward potassium currents were progressively decreased during repetitive depolarization (Fig. 1C), suggesting that progressive AP broadening most likely results from accumulation of inactivated potassium channels during bursts, as seen in other cell types (Geiger and Jonas, 2000; Kole et al., 2007). Occasionally, WT muscles also exhibited prolonged depolarizations (>150 ms), which are hereafter designated plateau potentials (PPs). PPs often occur at the end of an AP burst (Fig. 1A).

**Figure 1.**
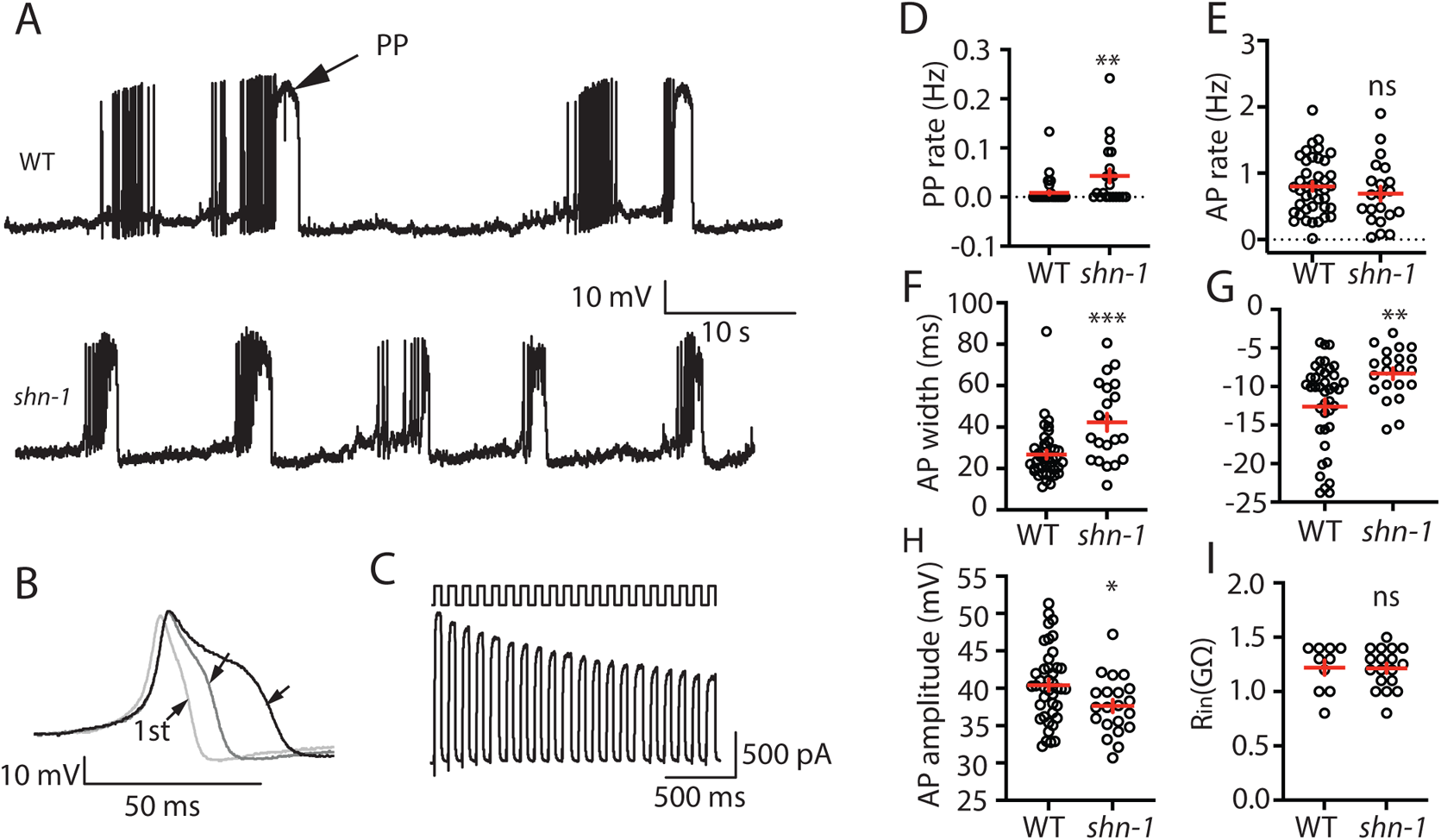
SHN-1 regulates muscle AP firing patterns. (A) Representative traces of spontaneous muscle APs are shown for WT and *shn-1(nu712)* null mutants. APs occur in bursts of ∼10 APs/ burst. Plateau potentials (PPs), defined as transients lasting >150 ms, are observed less frequently, often at the end of a burst. (B) APs become progressively longer during bursts. Successive APs taken from a representative burst are shown. (C) Repetitive depolarization to +30 mV leads to a progressive decrease in potassium currents. A representative recording from a WT animal is shown. This likely results from an accumulation of inactivated potassium channels during repetitive stimulation. (D-I) Mean PP rate (D), AP rate (E), AP width (F), RMP (G), AP amplitude (H), and input resistance (Rin, I) are compared in WT and *shn-1* null mutants. All *shn-1* data were obtained from *shn-1(nu712)* except for Rin (I), which were from *shn-1(tm488)*. Values that differ significantly from wild type controls are indicated (ns, not significant; *, *p* <0.05; **, *p* <0.01; ***, *p* <0.001). Error bars indicate SEM.

In *shn-1* null mutants, PP frequency and AP widths were significantly increased, AP amplitudes were decreased, resting membrane potential (RMP) was depolarized, while AP frequency and input resistance were unaltered (Fig. 1D-I). Similar increases in AP widths and PP frequency were observed in three, independently derived *shn-1* null mutants (*nu712*, *nu652*, and *tm488*) (Table 1). Single cell RNA sequencing studies suggest that SHN-1 is expressed in muscles, neurons, glia, and epithelial cells (Cao et al., 2017; Packer et al., 2019), consistent with the broad expression of split GFP tagged *shn-1(nu600* GFP_11_) (Fig. 1 supplement 1). To determine if SHN-1 functions in body muscles to control AP duration, we edited the endogenous *shn-1* locus to construct alleles that are either inactivated (*nu697*) or rescued (*nu652*) by the CRE recombinase (Fig. 2A). Using these alleles, we found that AP widths and PP frequency were increased in *shn-1(*muscle Knockout, KO) and that this defect was eliminated in *shn-1(*muscle rescue) (Fig. 2B-D). By contrast, *shn-1* knockout in neurons had no effect on PP rate or AP widths (Fig. 2C-D). Because CRE expression in muscles produced opposite changes in AP firing patterns in strains containing the *shn-1 nu697* and *nu652* alleles, these results are unlikely to be caused by toxicity associated with CRE expression (Speed et al., 2019). Collectively, these results suggest that SHN-1 acts in body muscles to control AP duration.

**Figure 2.**
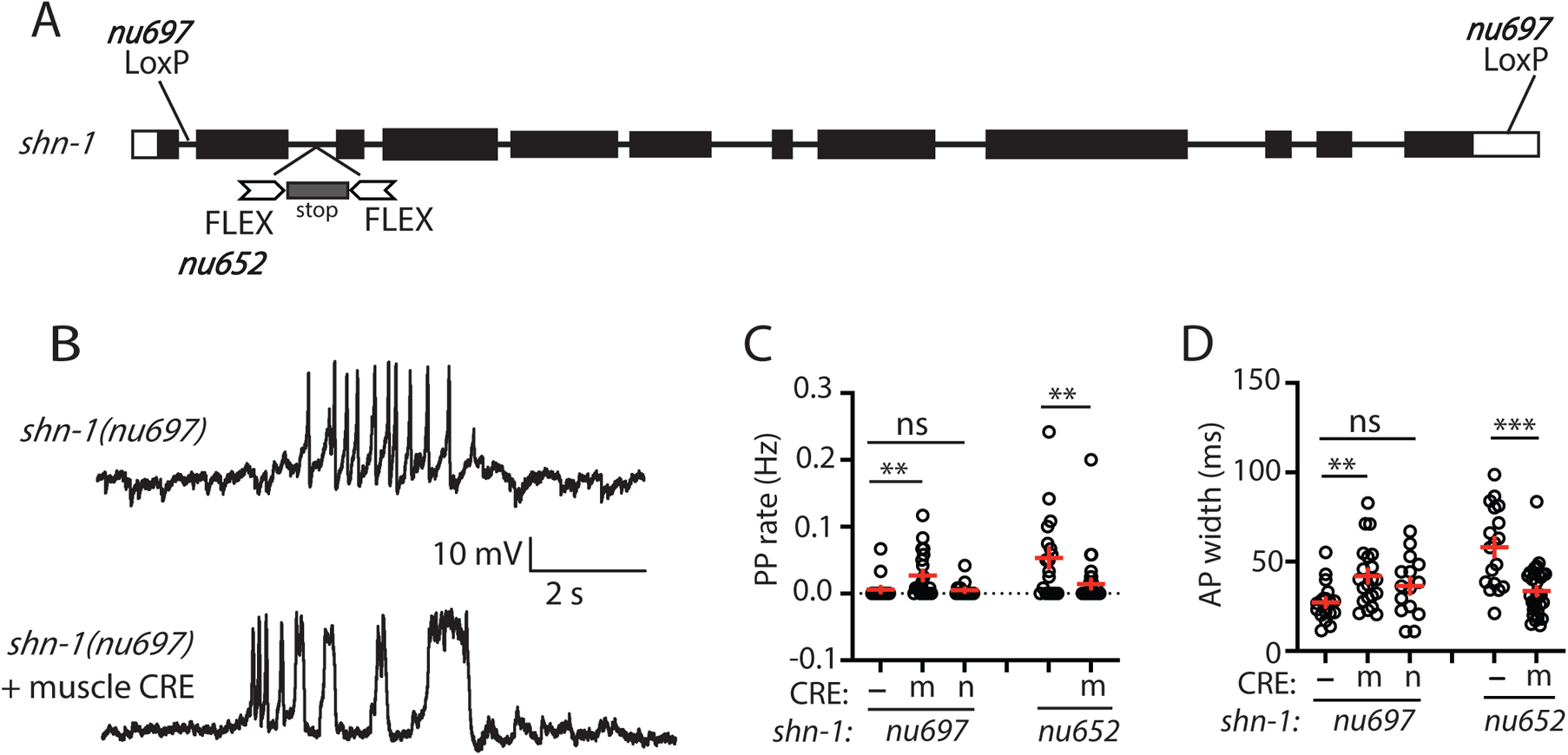
SHN-1 acts in muscles to control AP duration. (A) A schematic of the *shn-1* locus is shown. Open boxes indicate UTRs, black boxes indicate coding regions. Recombination sites mediating CRE induced deletions (LoxP) and inversions (FLEX) are indicated. The *shn-1(nu697)* allele allows CRE induced *shn-1* knockouts while *shn-1(nu652)* allows CRE induced *shn-1* rescue. In *shn-1(nu652)*, an exon containing in frame stop codons was inserted into the second intron (in the “OFF” orientation). This stop exon is bounded by FLEX sites. (B) Representative traces of spontaneous muscle APs are shown in *shn-1(nu697)* with and without muscle CRE expression. Mean PP rate (C) and AP width (D) are compared in the indicated *shn-1* mutants without (-) and with CRE expression in muscles (m) or neurons (n). Values that differ significantly from wild type controls are indicated (ns, not significant; *, *p* <0.05; **, *p* <0.01; ***, *p* <0.001). Error bars indicate SEM.

**Table 1.**
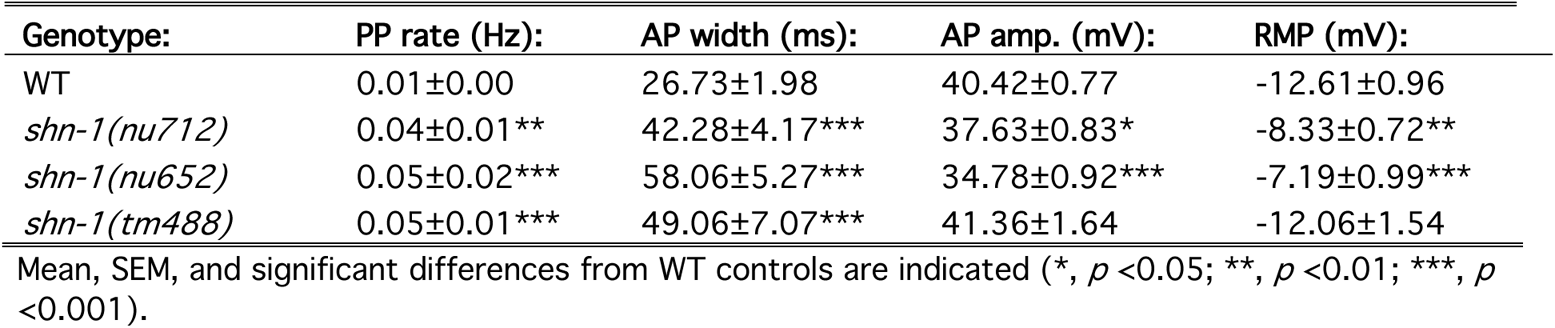
Comparison of shn-1 null alleles

### SHN-1 binding to EGL-19 promotes AP repolarization

Because SHN-1 has multiple binding partners, we sought to confirm that prolonged APs result from decreased SHN-1 binding to EGL-19. To address this question, we recorded APs in strains containing mutations that disrupt this interaction (Fig. 3A). PP frequency was significantly increased by a deletion removing the EGL-19 carboxy-terminal PDZ ligand [*egl-19(nu496 Δ*VTTL)] (Fig. 3B). AP widths were significantly increased by a deletion removing the SHN-1 PDZ domain [*shn-1(nu542 Δ*PDZ)] and by the *egl-19(nu496 Δ*VTTL) mutation (Fig. 3C). These results support the idea that SHN-1 binding to EGL-19/CaV1 accelerates AP repolarization.

**Figure 3.**
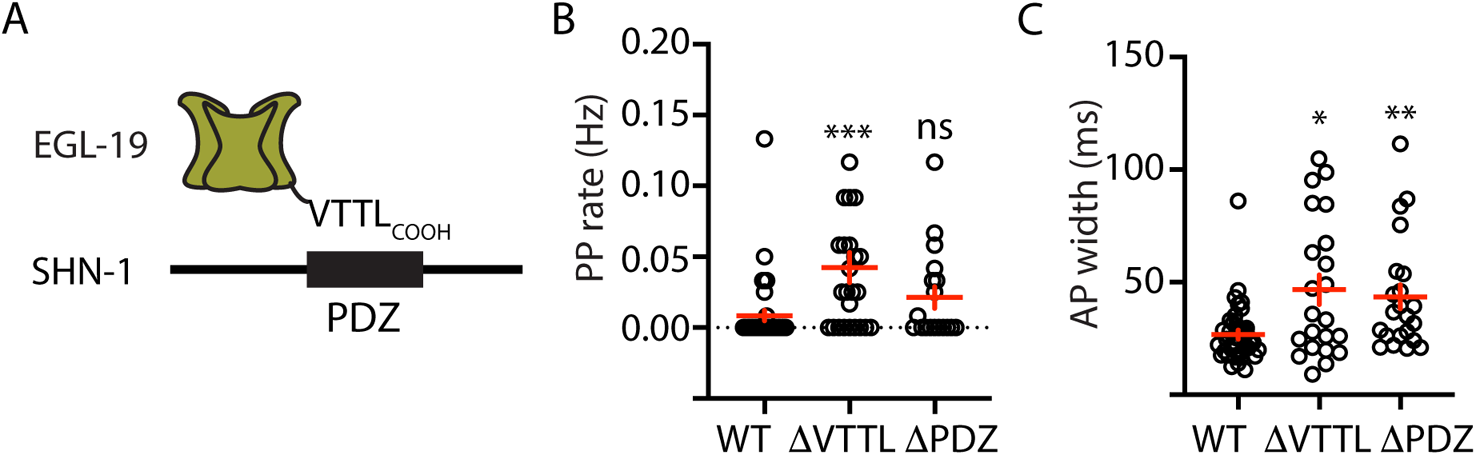
Mutations disrupting SHN-1 binding to EGL-19 increase AP duration. (A) A schematic illustrating the binding interaction between EGL-19’s c-terminus and SHN-1’s PDZ domain is shown. (B-C) Mean PP rate and AP width are compared in the indicated genotypes. Mutations deleting the SHN-1 PDZ domain (*nu542 Δ*PDZ) or those deleting EGL-19’s c-terminal PDZ ligand (*nu496 Δ*VTTL) were edited into the endogenous genes using CRISPR. These mutations significantly increased AP width, compared to WT controls. Values that differ significantly from wild type controls are indicated (ns, not significant; *, *p* <0.05; **, *p* <0.01; ***, *p* <0.001). Error bars indicate SEM.

### AP repolarization is controlled by SHK-1 KCNA and SLO-1/2 BK channels

To investigate how SHN-1 controls AP duration, we first asked which potassium channels promote repolarization following APs. Prior studies showed that voltage-activated potassium currents in body muscles are mediated by SHK-1/KCNA and BK channels (Gao and Zhen, 2011; Liu et al., 2011). SHK-1 channel function can be assessed in recordings using an internal solution containing low chloride levels (hereafter Ik_loCl_). Ik_loCl_ was nearly eliminated in *shk-1* single mutants (Fig. 4A-B). BK channel function can be assayed in recordings utilizing internal solutions with high chloride levels (hereafter Ik_hiCl_), which activates SLO-2 channels (Yuan et al., 2000). Ik_hiCl_ was ∼50% reduced in single mutants lacking either SLO-2 or SHK-1 and was eliminated in *slo-2; shk-1* double mutants (Fig. 4C-D). These results suggest that SHK-1/KCNA and SLO-2/BK are the primary channels promoting AP repolarization. Consistent with this idea, AP duration was significantly increased in mutants lacking SHK-1/KCNA (Fig. 4E-F), as previously reported (Gao and Zhen, 2011; Liu et al., 2011).

**Figure 4.**
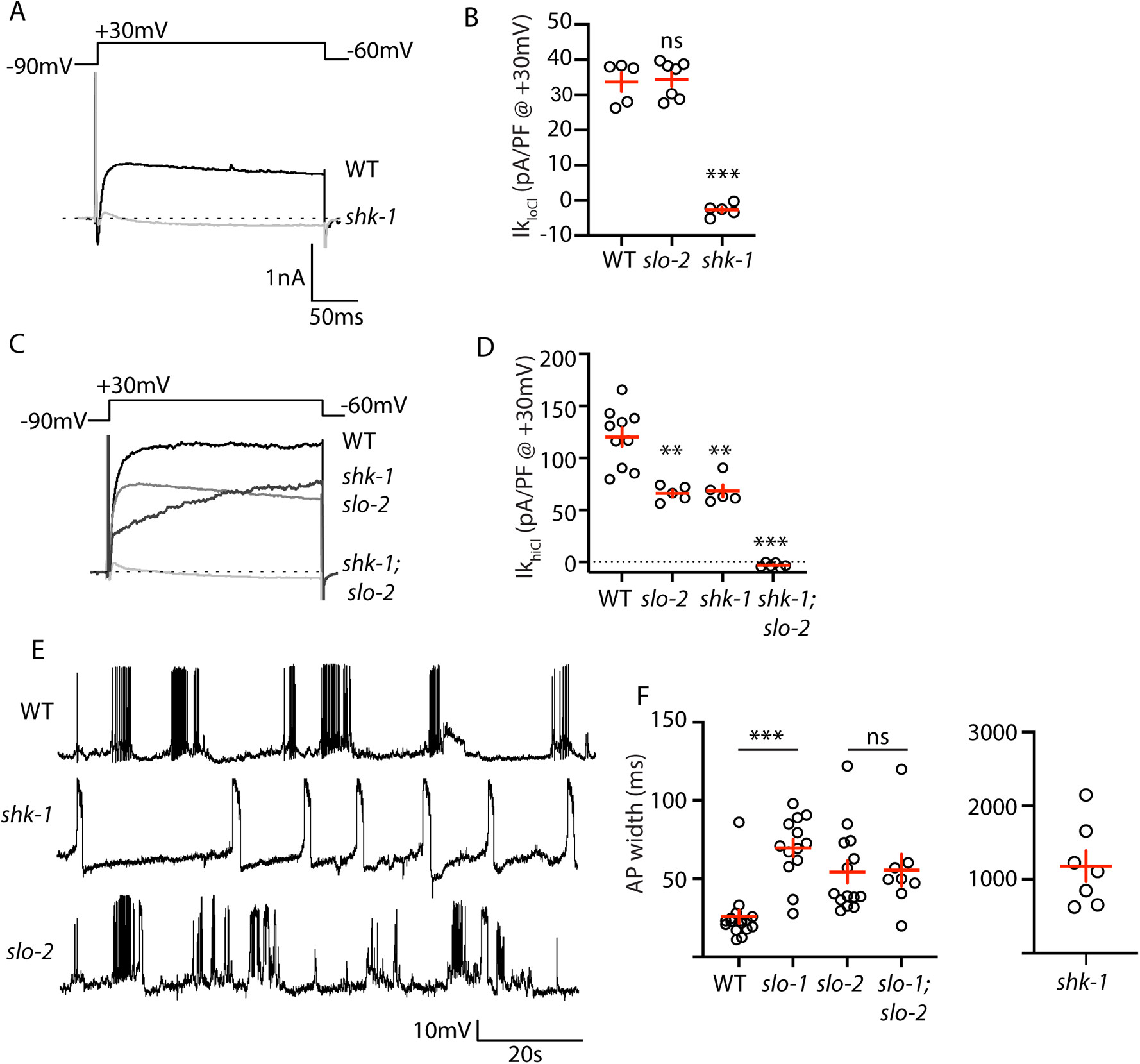
AP repolarization is mediated by SHK-1 and SLO-2 channels. (A-D) Muscle voltage activated potassium currents are mediated by SHK-1 and SLO-2. Voltage activated potassium currents were recorded using pipette solutions containing low (IkloCl, A-B) and high (IkhiCl, C-D) chloride concentrations. Representative traces (A,C) and mean current density (B,D) at +30 mV are shown. IkloCl is mediated by SHK-1 whereas SHK-1 and SLO-2 equally contribute to IkhiCl. (E-F) AP durations are significantly increased in mutants lacking SHK-1, SLO-1, and SLO-2 channels. The AP widths observed in *slo-1; slo-2* double mutants were not significantly different from those found in *slo-2* single mutants. Representative traces (E) and mean AP widths (F) are shown. Alleles used in this figure were: *shk-1(ok1581)*, *slo-1(js379)*, and *slo-2(nf100)*. Values that differ significantly from wild type controls are indicated (ns, not significant; *, *p* <0.05; **, *p* <0.01; ***, *p* <0.001). Error bars indicate SEM.

Contradictory results have been reported for AP firing patterns in *slo-1* and *slo-2* BK mutants (Gao and Zhen, 2011; Liu et al., 2011). These studies used intracellular solutions that alter BK channel function. In (Liu et al., 2011), an intracellular solution containing high chloride levels was used, thereby exaggerating SLO-2’s contribution to AP repolarization (Yuan et al., 2000). In (Gao and Zhen, 2011), an intracellular solution containing a fast calcium chelator (BAPTA) was used, which inhibits BK activation thereby minimizing their impact on APs. We re-investigated the effect of SLO channels on APs using intracellular solutions with low chloride and a slow calcium chelator (EGTA), finding that AP durations were increased to a similar extent in *slo-1* and *slo-2* single mutants (Fig. 4E-F). Taken together these results confirm that SHK-1 and SLO are the primary channels promoting AP repolarization in body muscles.

### SLO-1 and SLO-2 function together to promote AP repolarization

SLO-1 and SLO-2 subunits are co-expressed in muscles and could potentially co-assemble to form heteromeric channels. To determine if channels containing both SLO-1 and SLO-2 regulate AP repolarization, we analyzed AP widths in *slo-1; slo-2* double mutants. AP widths in *slo-1; slo-2* double mutants were not significantly different from those found in the single mutants (Fig. 4F). Because *slo-1* and *slo-2* mutations did not have additive effects on AP widths, these results support the idea that heteromeric SLO-1/2 channels mediate rapid repolarization of muscle APs.

We did several experiments to further test the idea that SLO-1 and SLO-2 function together in heteromeric channels. First, we recorded voltage activated potassium current in body muscles and found that Ik_hiCl_ was modestly reduced in *slo-1* mutants, was dramatically reduced in *slo-2* mutants, and was not further reduced in *slo-1; slo-2* double mutants (Fig. 5A-B). These results suggest that Ik_hiCl_ is mediated by heteromeric channels (containing both SLO-1 and SLO-2 subunits) and by SLO-2 homomers.

**Figure 5.**
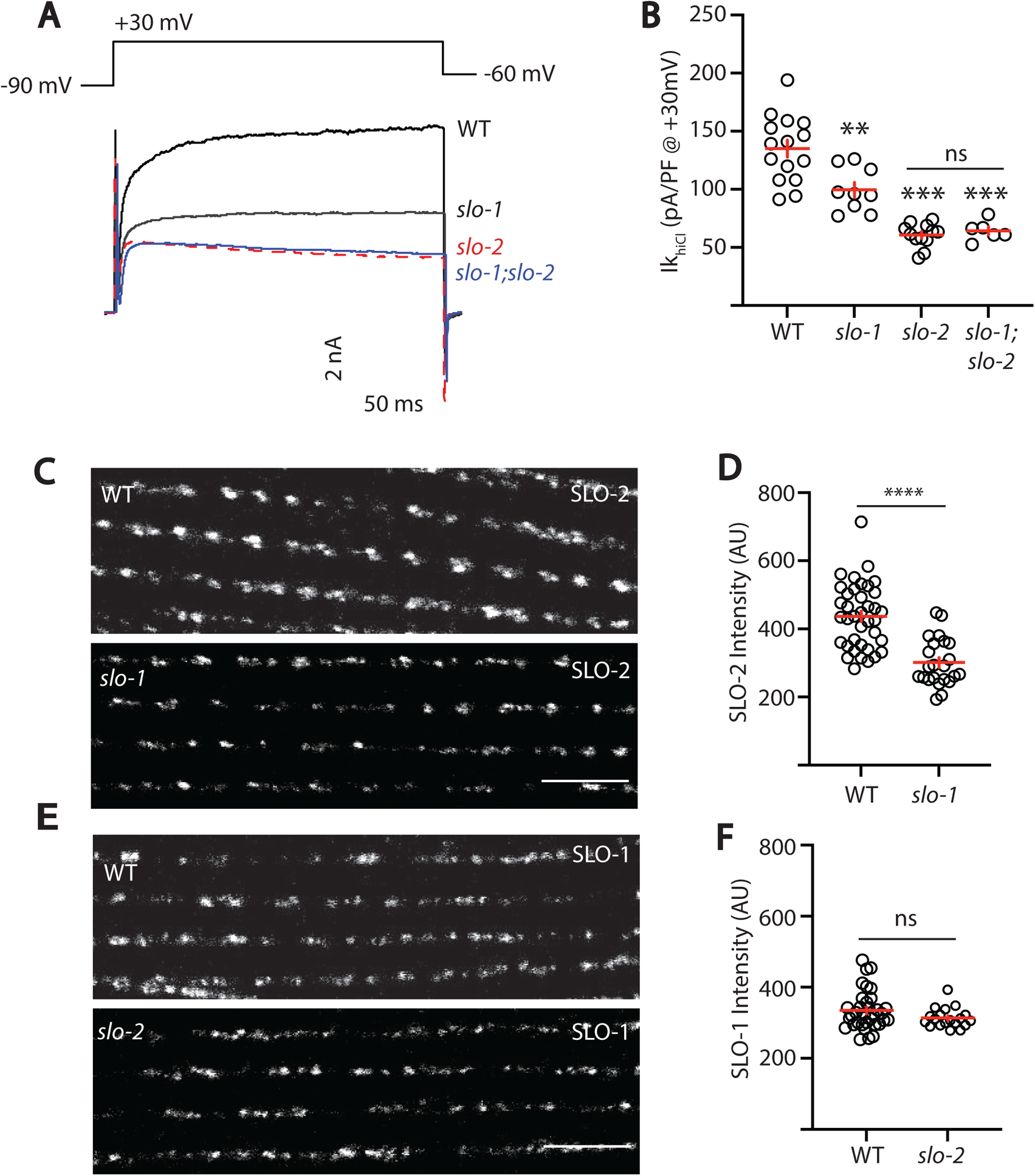
SLO-2 and SLO-1 function together in heteromeric channels. (A-B) IkhiCl was significantly decreased in *slo-1(js379)* and *slo-2(nf100)* single mutants but was not further decreased in *slo-1; slo-2* double mutants. Representative traces (A) and mean current density (B) at +30 mV are shown. (C-F) Expression of split GFP tagged SLO-2 (C-D) and SLO-1 (E-F) was analyzed in body muscles. CRISPR alleles were constructed adding 7 copies of GFP11 to the endogenous *slo-1* and *slo-2* genes (Table 2) and fluorescence was reconstituted by expressing GFP1-10 in body muscles. Controls showing that the GFP11 tags had no effect on AP width, RMP, and potassium currents are shown in Figure 5 supplement 1. Representative images (C and E) and mean puncta intensity (D and F) are shown. SLO-2 puncta intensity was significantly decreased in *slo-1(js379)* mutants. SLO-1 puncta intensity was unaltered in *slo-2(nf100)* mutants. Values that differ significantly from wild type controls are indicated (ns, not significant; *, *p* <0.05; **, *p* <0.01; ***, *p* <0.001). Error bars indicate SEM. Scale bar indicates 4 μm.

**Table 2.**
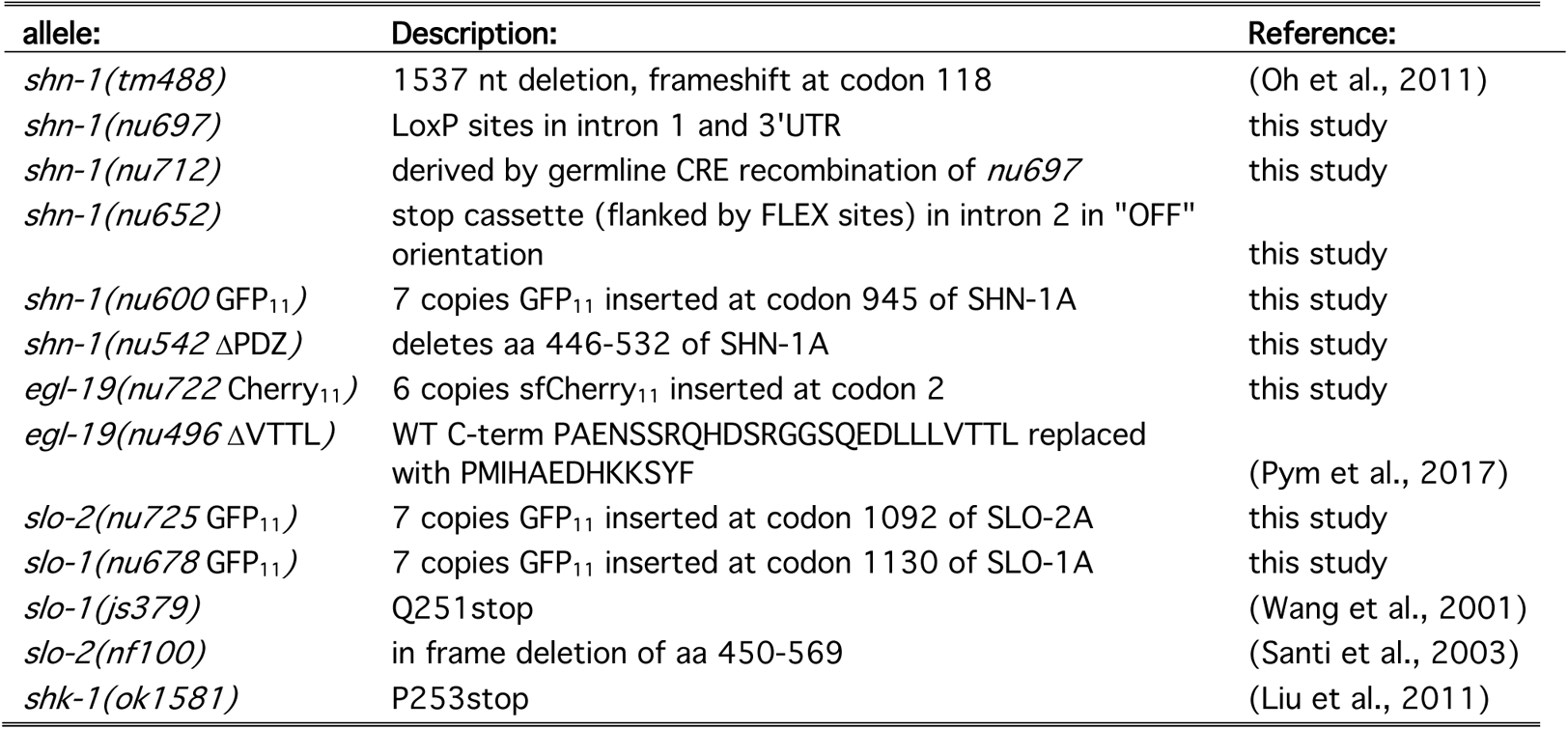
Alleles used in this study

As a final test of this idea, we asked if subcellular localization of SLO-1 and SLO-2 subunits requires expression of both subunits. For this analysis, endogenous SLO-1 and SLO-2 subunits were labelled with split GFP. Using CRISPR, we introduced the eleventh *β*-strand of GFP (GFP_11_) into the endogenous *slo-1* and *slo-2* genes and visualized their expression by expressing GFP_1-10_ in body muscles. Strains containing the GFP_11_ tagged alleles exhibited wild type Ik_hiCl_ currents, AP widths, and RMP, indicating that the tag did not interfere with SLO channel function (Fig. 5 supplement 1). Using these alleles, we find that SLO-2 puncta intensity was significantly reduced in *slo-1* null mutants, indicating that channels containing SLO-2 subunits require SLO-1 for their trafficking (Fig. 5C-D). By contrast, SLO-1 puncta intensity was unaffected in *slo-2* mutants, suggesting that BK channels lacking SLO-2 were trafficked normally (Fig. 5E-F). Collectively, these results suggest that rapid muscle repolarization following APs is mediated by SLO-1/2 heteromeric channels. Two prior studies also suggested that SLO subunits form heteromeric channels when heterologously expressed in *Xenopus* oocytes. SLO-1 currents were inhibited by a dominant-negative SLO-2 construct (Yuan et al., 2000). Similarly, mammalian SLO2 subunits (KCNT1 and 2) co-assemble to form heteromeric channels (Chen et al., 2009). Our results suggest that endogenously expressed SLO subunits also form heteromeric channels in native tissues.

### SHN-1 controls AP duration through BK channels

SHN-1’s impact on AP duration could be mediated by changes in either SHK-1 or SLO channels. To determine if SHN-1 acts through SLO channels, we asked if *slo-2* mutations block SHN-1’s effects on AP widths. Consistent with this idea, AP widths in *slo-2* single mutants were not significantly different from those in *slo-2* double mutants containing *shn-1(nu712* null), *shn-1(nu542 Δ*PDZ), or *egl-19(nu496 Δ*VTTL) mutations (Fig. 6A). These results suggest that SHN-1 controls AP duration by regulating SLO-2 channels.

**Figure 6.**
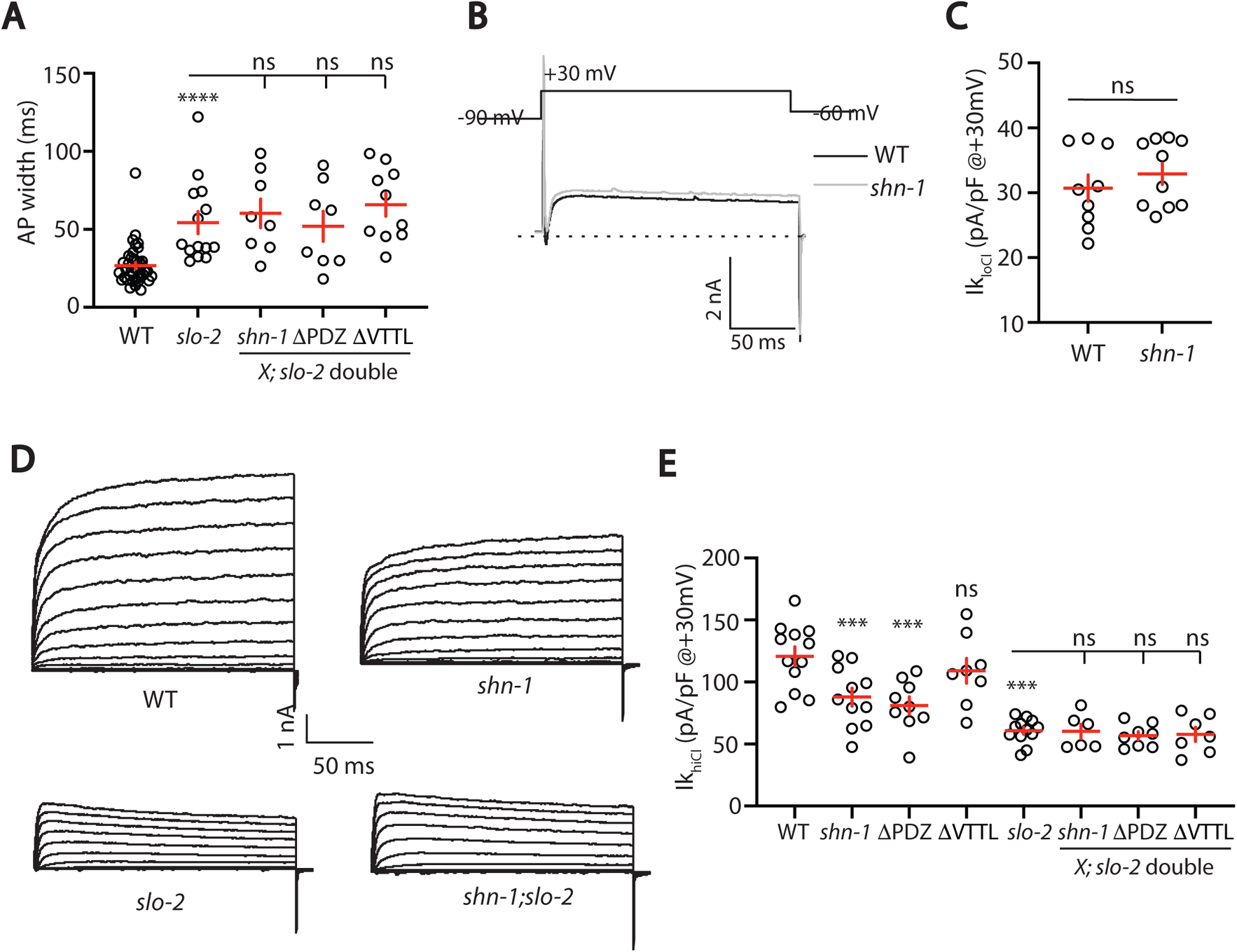
SHN-1 controls AP width by regulating SLO-2. (A) A *slo-2* null mutation blocks the effect of SHN-1 on AP width. Mean AP widths in *slo-2(nf100)* double mutants containing *shn-1(nu712* null*), shn-1(nu542 Δ*PDZ), and *egl-19(nu496 Δ*VTTL) mutations were not significantly different from those in *slo-2* single mutants. (B-C) IkloCl currents were unaltered in *shn-1(nu712* null) mutants. Representative traces (B) and mean current density at +30 mV (C) are shown. These results show that the function of SHK-1 channels was unaffected in *shn-1* mutants. (D-E) IkhiCl currents were significantly smaller in *shn-1(nu712* null) and *shn-1(nu542 Δ*PDZ) mutants but were unaffected in *egl-19(nu496 Δ*VTTL) mutants. The effect of *shn-1* mutations on IkhiCl was eliminated in double mutants lacking SLO-2, indicating that the SHN-1 sensitive potassium current is mediated by SLO-2. IkhiCl currents were recorded from adult body wall muscles of the indicated genotypes at holding potentials of −60 to +60 mV. Representative traces (D) and mean current density at +30 mV (E) are shown. Values that differ significantly from wild type controls are indicated (ns, not significant; *, *p* <0.05; **, *p* <0.01; ***, *p* <0.001). Error bars indicate SEM.

To confirm that SHN-1 regulates SLO-2 channels, we analyzed potassium currents in *shn-1* mutants. A *shn-1* null mutation had no effect on Ik_loCl_ currents, indicating that SHK-1 function was unaffected (Fig. 6B-C). By contrast, Ik_hiCl_ was ∼30% reduced in *shn-1* null mutants, ∼50% reduced in *slo-2* mutants, and was not further reduced in *shn-1; slo-2* double mutants (Fig. 6D-E). Lack of additivity in *shn-1; slo-2* double mutants suggests that the SHN-1 sensitive potassium current was mediated by SLO-2. A similar decrease in Ik_hiCl_ current was observed in *shn-1(nu542 Δ*PDZ) mutants (Fig. 6E). Ik_hiCl_ current was unaltered in *egl-19(nu496 Δ*VTTL) mutants, implying that SHN-1 binding to EGL-19’s carboxy terminus is not required for SLO-2 current (Fig. 6E). Thus, *shn-1* inactivation decreased SLO-1/2 BK current but had little or no effect on SHK-1 KCNA current; consequently, SHN-1 regulates AP widths by promoting activation of SLO-1/2 channels.

### SHN-1 promotes nanodomain coupling of SLO-2 with EGL-19/CaV1 channels

BK channels bind calcium with affinities ranging from 1-10 μM (Contreras et al., 2013). As a result of this calcium dependence, BK channels have very low open probability at resting cytoplasmic calcium levels (∼100 nM) and efficient BK activation typically requires close spatial coupling to calcium channels (Barrett et al., 1982). We next asked if body muscle BK channels are functionally coupled to EGL-19/CaV1 channels. Consistent with this idea, Ik_hiCl_ current was significantly decreased by nemadipine, an EGL-19/CaV1 antagonist (Fig. 7A-B) (Kwok et al., 2008). The inhibitory effect of nemadipine on Ik_hiCl_ was eliminated in *slo-2* mutants (Fig. 7A-B), suggesting that the nemadipine sensitive potassium current was mediated by SLO-2.

**Figure 7.**
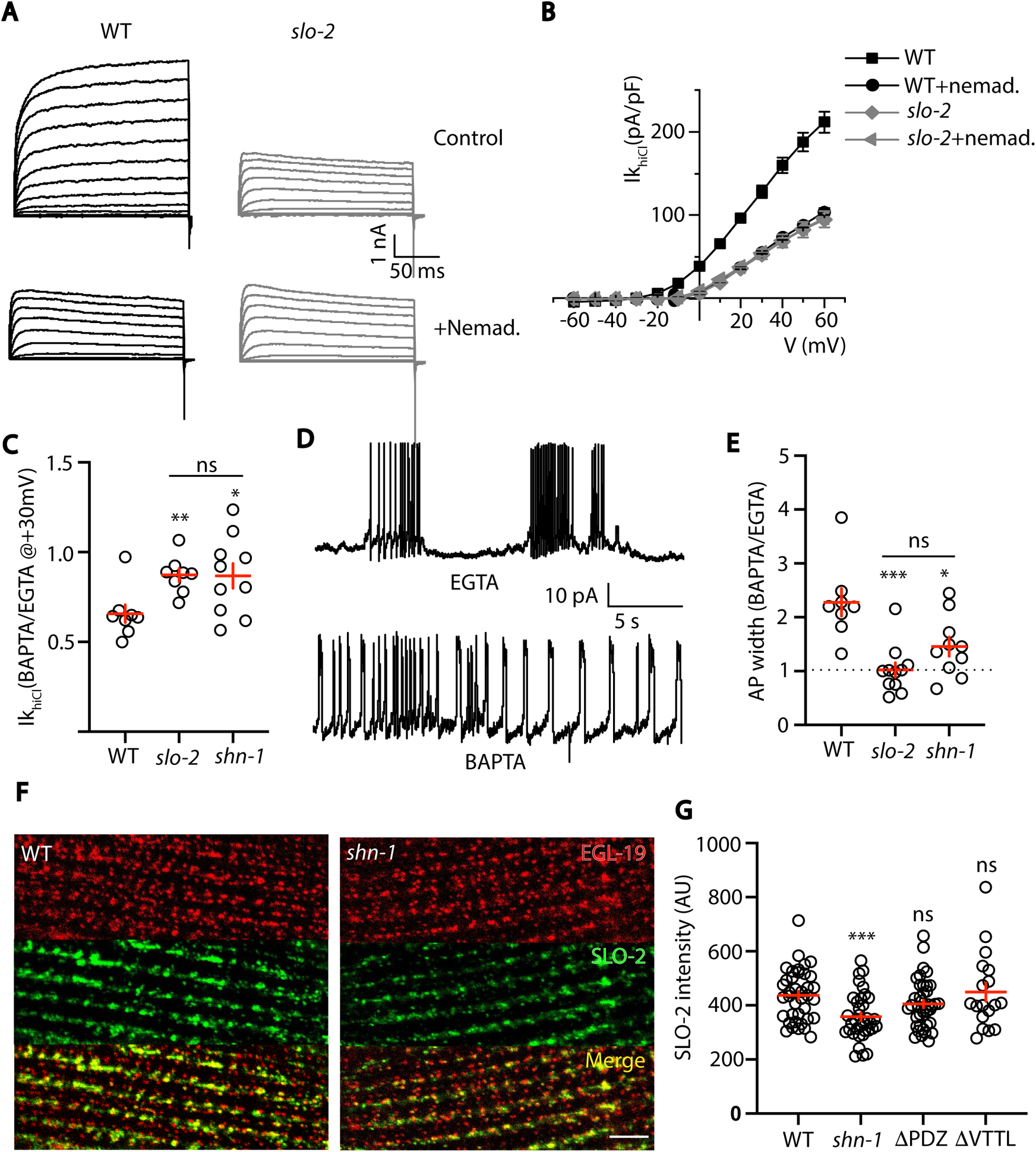
SHN-1 promotes EGL-19 to SLO-2 nanodomain coupling. (A-B) SLO-2 activation is functionally coupled to EGL-19. IkhiCl was significantly reduced by nemadapine (an EGL-19 antagonist). This inhibitory effect of nemadopine on IkhiCl was eliminated in *slo-2(nf100)* mutants, indicating that the nemadapine sensitive current is mediated by SLO-2. IkhiCl currents were recorded from adult body wall muscles of the indicated genotypes at holding potentials of −60 to +60 mV. Representative IkhiCl traces (A) and mean current density as a function of membrane potential (B) are shown. (C) SLO-2 activation requires nanodomain coupling to EGL-19. IkhiCl currents recorded in BAPTA are significantly smaller than those in EGTA. The inhibitory effect of BAPTA was reduced in *shn-1(nu712* null) mutants and was eliminated in *slo-2(nf100)* mutants, indicating that the BAPTA sensitive current is mediated by SLO-2. The ratio of IkhiCl current density at +30 mV recorded in BAPTA to the mean current density recorded in EGTA is plotted for the indicated genotypes. (D-E) AP repolarization is mediated by nanodomain activation of SLO-2. AP widths recorded in solutions containing BAPTA are wider than those recorded in EGTA. The effect of BAPTA on AP widths was reduced in *shn-1(nu712* null) mutants and was eliminated in *slo-2(nf100)* mutants, indicating that BAPTA’s effect is mediated by SLO-2. Representative traces of WT muscle APs recorded in EGTA and BAPTA are shown (D). The ratio of AP widths recorded in BAPTA to the mean AP widths recorded in EGTA is plotted for the indicated genotypes (E). (F-G) SLO-2(nu725 GFP11) is partially co-localized with EGL-19(nu722 Cherry11) in body muscles. GFP and Cherry fluorescence were reconstituted with GFP1-10 and Cherry1-10 expressed in body muscles. SLO-2 puncta intensity was significantly reduced in *shn-1(nu712* null) mutants but was unaffected in *shn-1(nu542 Δ*PDZ) and *egl-19(nu496 Δ*VTTL) mutants. Representative images (F) and mean SLO-2 puncta intensity (G) are shown. Values that differ significantly from wild type controls are indicated (ns, not significant; *, *p* <0.05; **, *p* <0.01; ***, *p* <0.001). Error bars indicate SEM. Scale bar indicates 4 μm.

Is EGL-19 coupling to SLO-2 mediated by nanodomain signaling? To test this idea, we compared Ik_hiCl_ and AP widths recorded with intracellular solutions containing fast (BAPTA) and slow (EGTA) calcium chelators (Fig. 7C-E). We found that Ik_hiCl_ recorded with BAPTA was significantly smaller than that recorded with EGTA (Fig. 7C). Similarly, AP widths recorded with BAPTA were significantly longer than those recorded with EGTA (Fig. 7D-E). The effect of BAPTA on Ik_hiCl_ and AP widths was eliminated in *slo-2* mutants (Fig. 7C,E), suggesting that the BAPTA sensitive potassium current is mediated by SLO-2. BAPTA’s effect on Ik_hiCl_ and AP widths was reduced but not eliminated in *shn-1* mutants (Fig. 7C,E), consistent with the partial loss of SLO-2 current in these mutants (Fig. 6D-E). Taken together, these results suggest that SHN-1 promotes SLO-2 nanodomain coupling to EGL-19/CaV1.

If BK channels are functionally coupled to EGL-19/CaV1, these channels should be co-localized. Endogenous SLO-2 channels (tagged with GFP11) were distributed in a punctate pattern on the muscle surface. A subset of the SLO-2 puncta co-localized with EGL-19/CaV1 channels (tagged with Cherry11), suggesting that EGL-19 nanocomplexes are heterogeneous (Fig. 7F). SLO-2 puncta intensity was significantly reduced in *shn-1* null mutants (Fig. 7F-G), consistent with the decreased SLO-2 current observed in these mutants. By contrast, SLO-2 puncta intensity was unaltered in *shn-1*(*nu542 Δ*PDZ) and *egl-19*(*nu496 Δ*VTTL) mutants (Fig. 7G), in which SHN-1 binding to EGL-19’s c-terminus is disrupted (Pym et al., 2017). SLO-1 puncta intensity was unaltered in *shn-1* null mutants, indicating that BK channels lacking SLO-2 were trafficked normally (Fig. 7 supplement 1). Collectively, these results suggest that SHN-1 stabilizes SLO-2 clusters in the plasma membrane and promotes activation of heteromeric SLO-1/2 channels by nearby calcium channels.

### EGL-19 to SLO-2 coupling is sensitive to *shn-1* gene dose

Deletion and duplication of human shank genes are both associated with ASD, schizophrenia, and mania (Bonaglia et al., 2006; Durand et al., 2007; Failla et al., 2007; Gauthier et al., 2010; Han et al., 2013). These results suggest that Shank phenotypes relevant to psychiatric disorders should exhibit a similar sensitivity to Shank copy number. For this reason, we analyzed the effect of *shn-1* gene dosage on Ik_hiCl_ and AP duration (Fig. 8). We analyzed animals with 1 (*nu712/+* heterozygotes), 2 (WT), and 4 (WT+2 single copy *shn-1* transgenes) copies of *shn-1*. Compared to wild type controls, muscle Ik_hiCl_ was significantly decreased while AP duration was significantly increased in animals containing 1 and 4 copies of *shn-1* (Fig. 8). Thus, increased and decreased *shn-1* gene dosage produced similar defects in AP duration and SLO-2 current.

**Figure 8.**
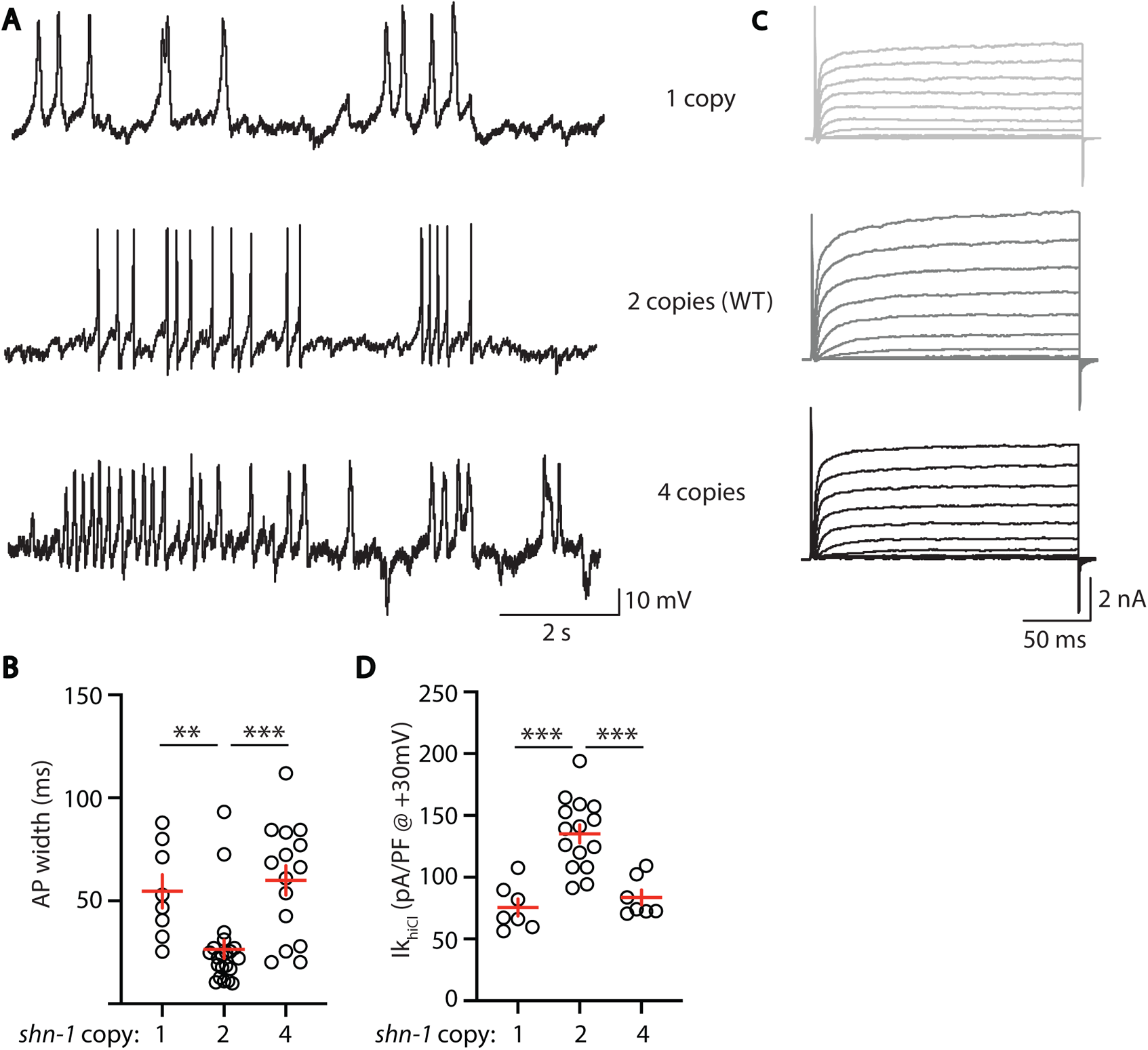
AP width and SLO-2 current are sensitive to *shn-1* gene dosage. The effect of *shn-1* gene dosage on AP widths (A-B) and IkhiCl current (C-D) was analyzed. Ik_hiCl_ was significantly decreased while AP duration was significantly increased in animals containing 1 and 4 copies of *shn-1* compared to WT controls (i.e. 2 copies). The following genotypes were analyzed: 1 copy of *shn-1* [*shn-1(nu712)*/+ heterozygotes], 2 copies of *shn-1* (WT) and 4 copies of *shn-1* (*nuSi26* homozygotes in wild-type). IkhiCl currents were recorded from adult body wall muscles of the indicated genotypes at holding potentials of −60 to +60 mV. Representative traces (A,C), mean AP width (B), and mean IkhiCl current density at +30 mV (D) are shown. Significant differences are indicated (ns, not significant; *, *p* <0.05; **, *p* <0.01; ***, *p* <0.001). Error bars indicate SEM.

### SHN-1 regulates BK channel activation and neurotransmitter release in motor neurons

Thus far, our results suggest that SHN-1 promotes EGL-19 to SLO-2 coupling in muscles. We next asked if SHN-1 also promotes coupling in motor neurons. To test this idea, we analyzed Ik_hiCl_ in cholinergic motor neurons and found that it was significantly reduced in *shn-1* null mutants (Fig. 9A-C). The *slo-2* and *shn-1* mutations did not have additive effects on Ik_hiCl_ in double mutants, suggesting that the *shn-1* mutation selectively decreases SLO-2 current in motor neurons (Fig. 9A-C). Consistent with decreased SLO-2 currents, we observed a corresponding decrease in axonal SLO-2(*nu725* GFP_11_) puncta fluorescence in *shn-1* mutant motor neurons (Fig. 9D-E). Thus, our results suggest that SHN-1 promotes EGL-19/CaV1 to SLO-2 coupling in both body muscles and cholinergic motor neurons.

**Figure 9.**
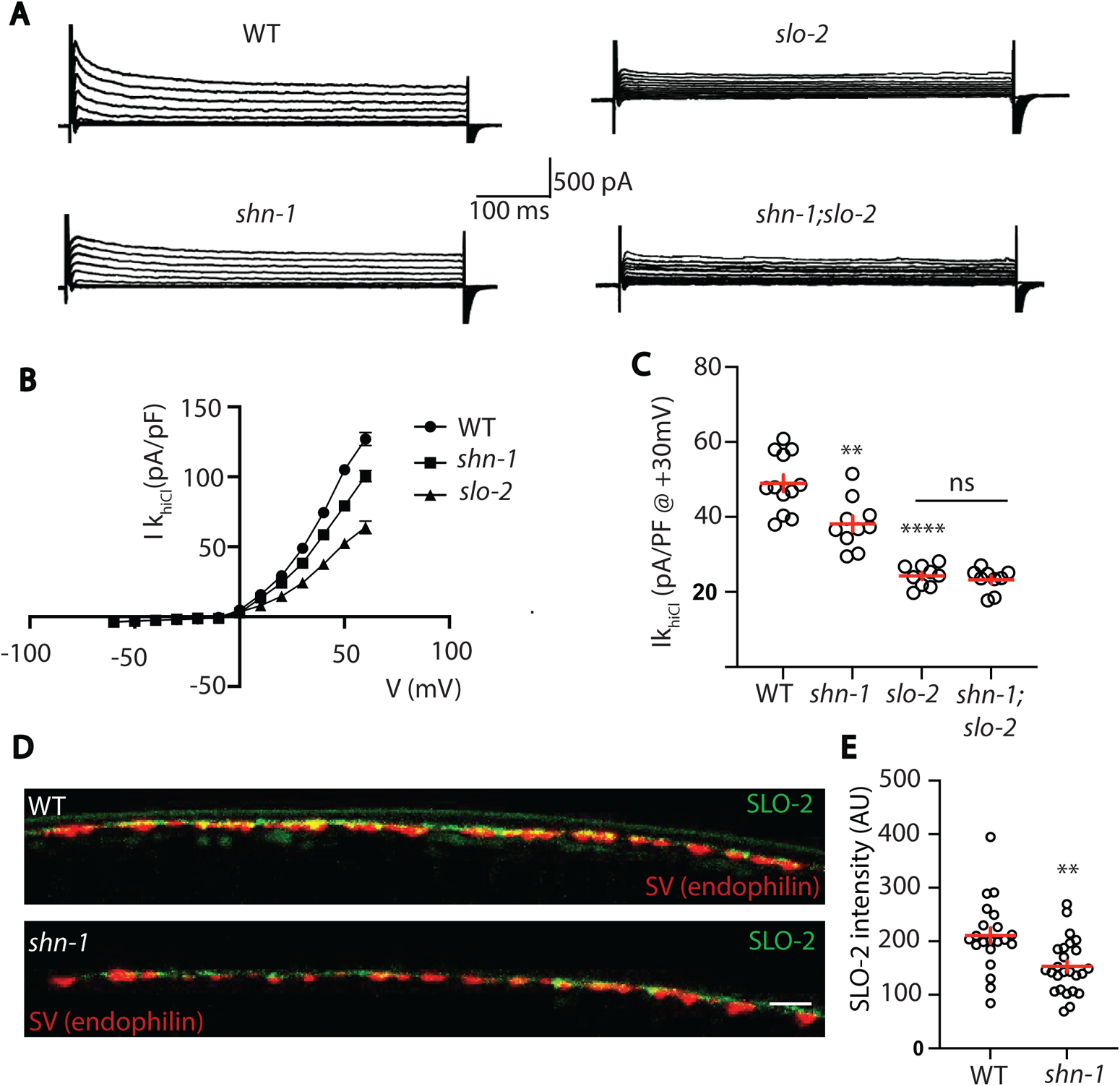
SHN-1 controls SLO-2 currents in motor neurons. (A-B) IkhiCl currents in cholinergic motor neurons were significantly decreased in *shn-1(nu712* null) mutants. IkhiCl currents were recorded from adult cholinergic motor neurons of the indicated genotypes at holding potentials of −60 to +60 mV. Representative traces (A), mean current density as a function of membrane potential (B), and mean current density at +30 mV (C) are shown. (D-E) SLO-2 puncta intensity in motor neuron axons was significantly decreased in *shn-1(nu712* null) mutants. Representative images of SLO-2(*nu725* GFP11) and a synaptic vesicle marker [UNC-57/Endophilin(mCherry)] in dorsal cord axons of DA/DB motor neurons are shown (D). GFP11 fluorescence was reconstituted with GFP1-10 expressed in DA/DB motor neurons (using the *unc-129* promoter). Mean SLO-2 puncta intensity in axons is plotted (E). Values that differ significantly from wild type controls are indicated (ns, not significant; *, *p* <0.05; **, *p* <0.01; ***, *p* <0.001). Error bars indicate SEM. Scale bar indicates 2 μm.

## Discussion

Our results lead to six principal conclusions. First, we show that SHN-1 acts cell autonomously in muscles to promote rapid repolarization of APs. Second, heteromeric BK channels containing both SLO-1 and SLO-2 subunits promote AP repolarization. Third, SHN-1 limits AP duration by promoting BK channel activation. Fourth, *shn-1* mutants have decreased SLO-2 channel clustering and decreased SLO-2 currents. Fifth, increased and decreased SHN-1 gene dosage produce similar defects in AP durations and SLO-2 currents. And sixth, SHN-1 also promotes SLO-2 activation in motor neurons. Below we discuss the significance of these findings.

### Shank as a regulator of ion channel density

Several recent studies suggest that an important function of Shank proteins is to regulate ion channel density and localization. Mutations inactivating Shank have been shown to decrease AMPA and NMDA receptor abundance and post-synaptic currents (Peca et al., 2011; Won et al., 2012), HCN channels (Yi et al., 2016; Zhu et al., 2018), TRPV channels (Han et al., 2016), and voltage activated CaV1 calcium channels (Pym et al., 2017; Wang et al., 2017). Here we show that Shank also regulates BK channel densities in *C. elegans* muscles and motor neurons. Collectively, these studies suggest that Shank proteins have the capacity to control localization of many ion channels, thereby shaping neuron and muscle excitability.

Shank regulation of BK channels could have broad effects on neuron and muscle function. In neurons, BK channels are functionally coupled to CaV channels in the soma and dendrites, thereby regulating AP firing patterns and somatodendritic calcium transients (Golding et al., 1999; Storm, 1987). In pre-synaptic terminals, BK channels limit the duration of calcium influx during APs, thereby decreasing neurotransmitter release (Griguoli et al., 2016; Yazejian et al., 2000). In muscles, BK channels regulate AP firing patterns, calcium influx during APs, and muscle contraction (Dopico et al., 2018; Latorre et al., 2017). Thus, Shank mutations could broadly alter neuron and muscle function via changes in CaV-BK coupling. It will be very interesting to determine if this new function for Shank is conserved in other animals, including humans.

### SHN-1 promotes nanodomain coupling of CaV1 and BK channels

BK channel activation requires tight coupling to CaV channels (Berkefeld et al., 2006). Our results suggest that SHN-1 promotes CaV1-BK nanodomain coupling. APs were prolonged in *shn-1* (null), *shn-1(Δ*PDZ), and *egl-19(Δ*VTTL) mutants and in all cases these defects were eliminated in double mutants lacking SLO-2. Interestingly, although all impair SLO-2 mediated AP repolarization, these mutations had distinct effects on SLO-2 channels. SLO-2 currents were reduced in *shn-1* (null) and *shn-1(Δ*PDZ) mutants but were unaffected in *egl-19(Δ*VTTL) mutants. SLO-2 puncta intensity was decreased in *shn-1*(null) mutants but was unaffected in *shn-1(Δ*PDZ) and *egl-19(Δ*VTTL) mutants.

These differences suggest that these mutants comprise an allelic series for EGL-19 to SLO-2 coupling defects in the following hierarchy *shn-1* (null)>*shn-1(Δ*PDZ)> *egl-19(Δ*VTTL) mutants. Based on these results, we propose that multiple protein interactions progressively tighten CaV-BK coupling. Specifically, we propose that: 1) multiple SHN-1 domains act together to promote SLO-2 coupling to EGL-19, accounting for the distinct phenotypes observed in *shn-1(*null) and *shn-1(Δ*PDZ) mutants; 2) SHN-1 promotes formation (or stability) of SLO-2 clusters in the plasma membrane, as indicated by decreased SLO-2 puncta intensity in *shn-1(*null) mutants; 3) beyond this trafficking function, SHN-1’s PDZ domain tightens SLO-2 coupling to nearby calcium channels, accounting for the smaller SLO-2 current but unaltered SLO-2 puncta intensity in *shn-1(Δ*PDZ) mutants; 4) SHN-1 PDZ binding to EGL-19’s c-terminus promotes rapid SLO-2 activation during APs, accounting for the increased AP width but unaltered SLO-2 current and puncta intensity in *egl-19(Δ*VTTL) mutants; and 5) SHN-1’s PDZ must bind multiple proteins (not just EGL-19) to promote SLO-2 activation, accounting for the different phenotypes found in in *shn-1(Δ*PDZ) and *egl-19(Δ*VTTL) mutants. Multivalent interactions between scaffolds and their client proteins may represent a general mechanism for promoting nanodomain signaling.

### Implications for understanding neurodevelopmental disorders

Shank3 deletions and duplications both confer risk for ASD and schizophrenia (Bonaglia et al., 2006; Durand et al., 2007; Failla et al., 2007; Gauthier et al., 2010; Han et al., 2013). It is currently unclear how opposite changes in Shank3 levels produce similar psychiatric phenotypes. Different (potentially opposite) biochemical defects arising from decreased and increased Shank dosage could produce similar psychiatric traits, perhaps by circuit level mechanisms (Antoine et al., 2019; Peixoto et al., 2016). For example, Shank duplications and hemizygosity could act in different cells or circuits to produce similar psychiatric traits. Our results provide support for a second possibility. We find that increased and decreased *shn-1* gene dosage produce similar cell autonomous CaV1-BK coupling defects. Two prior studies suggest that bidirectional changes in Shank produce similar defects in Wnt signaling and CaV1 current density (Harris et al., 2016; Pym et al., 2017). Collectively, these results suggest that some biochemical functions of Shank exhibit this unusual pattern of dose sensitivity and consequently could contribute to the pathophysiology of human Shankopathies (i.e. Shank3 mutations, CNVs, or PMS).

The role of human Shank in CaV1-BK coupling has not been tested. Nonetheless, it seems plausible that this new physiological function could contribute to neuropsychiatric or co-morbid phenotypes associated with human Shankopathies. Consistent with this idea, PMS and human KCNMA1/BK mutations are associated with several shared phenotypes including: autism, developmental delay, intellectual disability, hypotonia, seizures, and gastrointestinal defects (i.e. vomiting, constipation, or diarrhea) (Bailey et al., 2019; Laumonnier et al., 2006; Phelan and McDermid, 2012; Soorya et al., 2013; Witmer et al., 2019). Moreover, BK channels are regulated by two high confidence ASD genes (UBE3A and hnRNP U). BK channels are degraded by the ubiquitin ligase UBE3A (Sun et al., 2019), mutations in which cause Angelman’s syndrome. The RNA binding protein hnRNP U promotes translation of *slo-2* mRNA (Liu et al., 2018).

Collectively, these results support the idea that disrupted CaV1-BK channel coupling could play an important role in shankopathies and that BK channels may represent an important therapeutic target for treating these disorders.

### Experimental Procedures

*Strains:* Strain maintenance and genetic manipulation were performed as described (Brenner, 1974). Animals were cultivated at room temperature (∼22°C) on agar nematode growth media seeded with OP50 bacteria. Alleles used in this study are described in Table 2 and are identified in each figure legend. Transgenic animals were prepared by microinjection, and integrated transgenes were isolated following UV irradiation, as described (Dittman and Kaplan, 2006). Single copy transgenes were isolated by the MoSCI and miniMoS techniques (Frokjaer-Jensen et al., 2008; Frokjaer-Jensen et al., 2014).

### *shn-1* dosage experiments

Animals with different *shn-1* copy numbers were constructed as follows: 0 copies, *shn-1*(*nu712)* homozygotes; 1 copy, *unc-17::gfp (LX929)* males were crossed with *shn-1*(*nu712)* homozygotes and *gfp*-expressing hermaphrodites were analyzed; 2 copies, WT were analyzed; 4 copies, WT animals homozygous for the single copy transgene expressing SHN-1A in body muscles (*nuSi26*). The *nuSi26* transgene was described in our earlier study (Pym et al., 2017).

### CRISPR alleles

CRISPR alleles were isolated as described (Arribere et al., 2014). Briefly, we used *unc-58* as a co-CRISPR selection to identify edited animals. Animals were injected with two guide RNAs (gRNAs) and two repair templates, one introducing an *unc-58* gain of function mutation and a second modifying a gene of interest. Progeny exhibiting the *unc-58(gf)* uncoordinated phenotype were screened for successful editing of the second locus by PCR. Split GFP and split sfCherry constructs are described in (Feng et al., 2017). MiniMOS inserts in which Pmyo-3 drives expression of either GFP1-10 (nuTi144) or sfCherry1-10 SL2 GFP1-10 (nuTi458) were created.

Tissue specific *shn-1* knockout was performed by introducing LoxP sites into intron 1 and the 3’UTR of the endogenous locus, in *shn-1(nu697)*, and expressing CRE in muscles (*pat-10* promoter) or neurons (*sbt-1* promoter). Tissue specific *shn-1* rescue was performed by introducing a stop cassette into intron 2 of *shn-1* using CRISPR, creating the *shn-1(nu652)* allele. The stop cassette consists of a synthetic exon (containing a consensus splice acceptor sequence and stop codons in all reading frames) followed by a 3’ UTR and transcriptional terminator taken from the *flp-28* gene (the 564bp sequence just 3’ to the *flp-28* stop codon). The stop cassette is flanked by FLEX sites (which are modified loxP sites that mediate CRE induced inversions) (Schnutgen and Ghyselinck, 2007). In this manner, orientation of the stop cassette within the *shn-1* locus is controlled by CRE expression. Expression of *shn-1* is reduced when the stop cassette is in the OFF configuration (i.e. the same orientation as *shn-1*) but is unaffected in the ON configuration (opposite orientation). The endogenous *flp-28* gene is located in an intron of W07E11.1 (in the opposite orientation). Consequently, we reasoned that the *flp-28* transcriptional terminator would interfere with *shn-1* expression in an orientation selective manner. A similar strategy was previously described for conditional gene knockouts in *Drosophila* (Fisher et al., 2017).

### Fluorescence imaging

Confocal imaging was performed using a Nikon 60x objective (NA 1.45) on a Nikon AR1 confocal microscope. Worms were immobilized on 10% agarose pads with 0.3 µl of 0.1 µm diameter polystyrene microspheres (Polysciences 00876-15, 2.5% w/v suspension). Muscles just anterior to the vulva were imaged. For quantitation of florescence intensities, puncta were analyzed using Fiji.

### Electrophysiology

Whole-cell patch-clamp measurements were performed using a Axopatch 200B amplifier with pClamp 10 software (Molecular Devices). The data were sampled at 10 kHz and filtered at 5 kHz. All recordings were performed at room temperature (∼19-21 °C) *Muscle APs*-The bath solution contained (in mM): NaCl 140, KCl 5, CaCl2 5, MgCl2 5, dextrose 11 and HEPES 5 (pH 7.2, 320 mOsm). The pipette solution contained (in mM): Kgluconate 120, KOH 20, Tris 5, CaCl2 0.25, MgCl2 4, sucrose 36, EGTA 5 (or BAPTA 5), and Na2ATP 4 (pH 7.2, 323 mOsm). Spontaneous APs were recorded in current-clamp without current injection. Cell resistance (Rin) was measured following a 10 pA pulse injection. AP traces were analyzed in Matlab. APs were defined as depolarizations lasting <150 ms. PPs were defined as depolarizations lasting >150 ms. *K^+^ current recordings* - The bath solution contained (in mM): NaCl 140, KCl 5, CaCl2 5, MgCl2 5, dextrose 11 and HEPES 5 (pH 7.2, 320 mOsm). For Ik_loCl_ recordings, the pipette solution contained (in mM): Kgluconate 120, KOH 20, Tris 5, CaCl2 0.25, MgCl2 4, sucrose 36, EGTA 5 and Na2ATP 4 (pH 7.2, 323 mOsm). For Ik_hiCl_ recordings, the pipette solution contained (in mM): KCl 120, KOH 20, Tris 5, CaCl2 0.25, MgCl2 4, sucrose 36, EGTA 5 (or BAPTA 5), and Na2ATP 4 (pH 7.2, 323 mOsm). The voltage-clamp protocol consisted of −60mV for 50ms, −90mV for 50 ms, test voltage (from −60mV to +60mV) 150 ms. The repetitive stimulus protocol was −20mV for 50ms, +30mV for 50ms, which was repeated 20 times. In figures, we show outward currents evoked at +30 mV, which corresponds to the peak amplitude of muscle APs. In some recordings, EGL-19 channels were blocked by adding 5μM nemadipine to the pipette solution. Patch clamp recording of IkhiCl in ACh motor neurons was done using solutions described above for the muscle recordings. ACh neurons were identified for patching by expression of *unc-17* VAChT>GFP.

### Statistical methods

For normally distributed data, significant differences were assessed with unpaired t tests (for 2 groups) or one way ANOVA with post-hoc Dunn’s multiple comparisons test (for >2 groups). For non-normal data, differences were assessed by Mann-Whitney (2 groups) or Kruskal-Wallis test with post-hoc Dunn’s multiple comparisons test (>2 groups). Data graphing and statistics were performed in GraphPad Prism 9. No statistical method was used to select sample sizes. Data shown in each figure represent contemporaneous measurements from mutant and control animals over a period of 1-2 weeks. For electrophysiology, data points represent mean values for individual neuron or muscle recordings (which were considered biological replicates). For imaging studies, data points represent mean puncta fluorescence values in individual animals (which were considered biological replicates). All data obtained in each experiment were analyzed, without any exclusions.

## Acknowledgements

We thank the following for strains, advice, reagents, and comments on the manuscript: *C. elegans* stock center, S. Mitani, and members of the Kaplan lab. This work was supported by an NIH research grant to J.K. (NS32196).

---

**Figure 1, Supplement 1.**
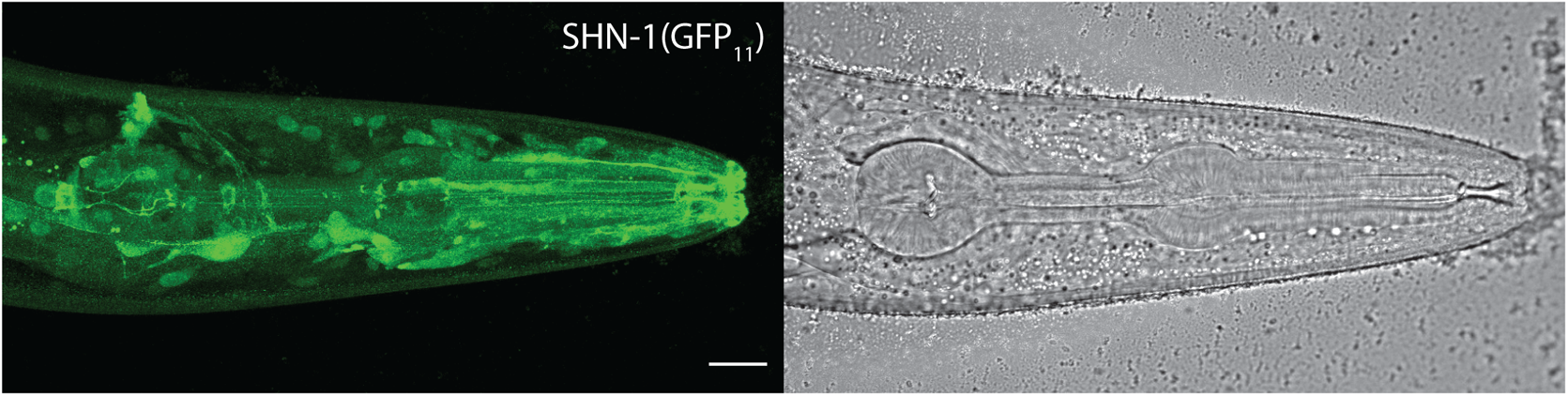
SHN-1 is expressed in many tissues. Endogenous SHN-1 is broadly expressed, including in neurons, muscles, skin, and glia. A representative image of reconstituted fluorescence produced by *shn-1(nu600* GFP11) and eft-3>GFP1-10 (left) and the corresponding bright field image (right) are shown. Scale bar indicates 14 μm.

**Figure 5. Supplement 1.**
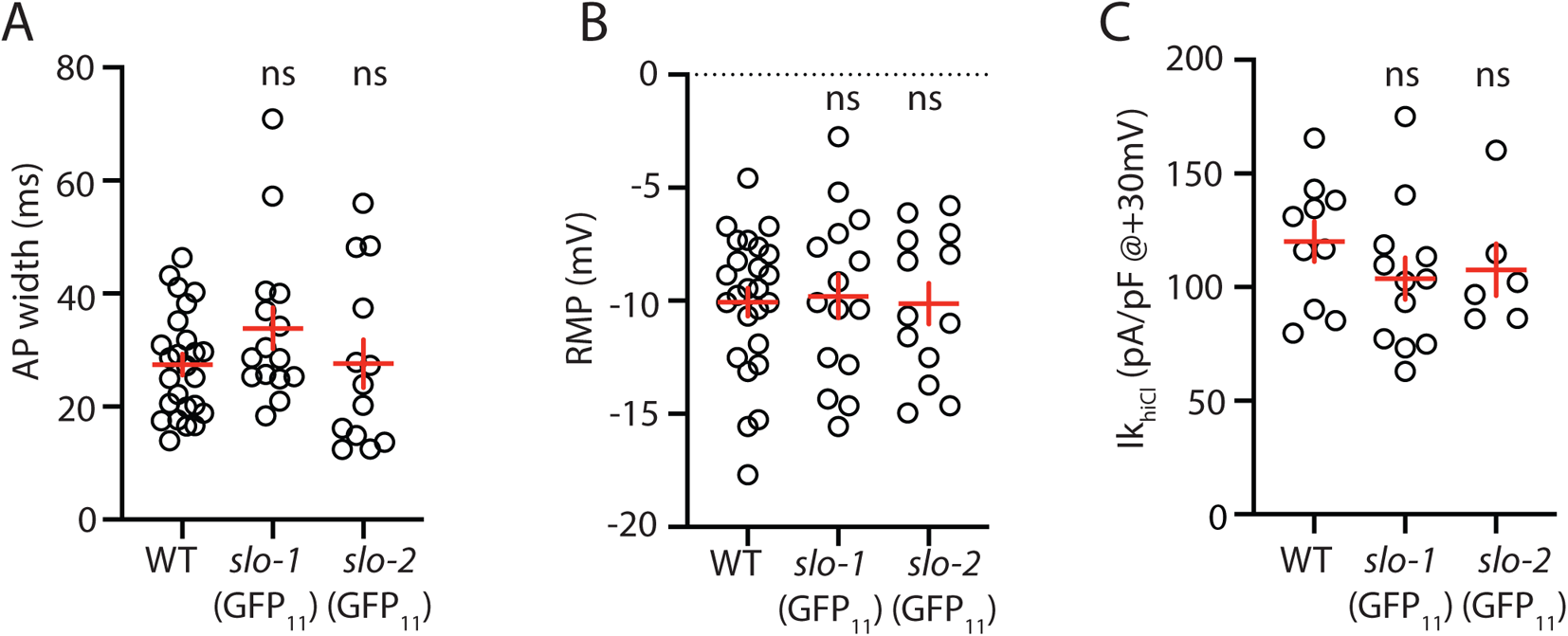
Analysis of GFP11 tagged *slo-1* and *slo-2* alleles. APs and potassium currents were analyzed in strains containing *slo-1(nu678* GFP11) and *slo-2(nu725* GFP11) together with the muscle>GFP1-10 transgene. Mean AP width (A), RMP (B), and IkhiCl current density (C) were not significantly different from WT controls. Error bars indicate SEM.

**Figure 7. Supplement 1.**
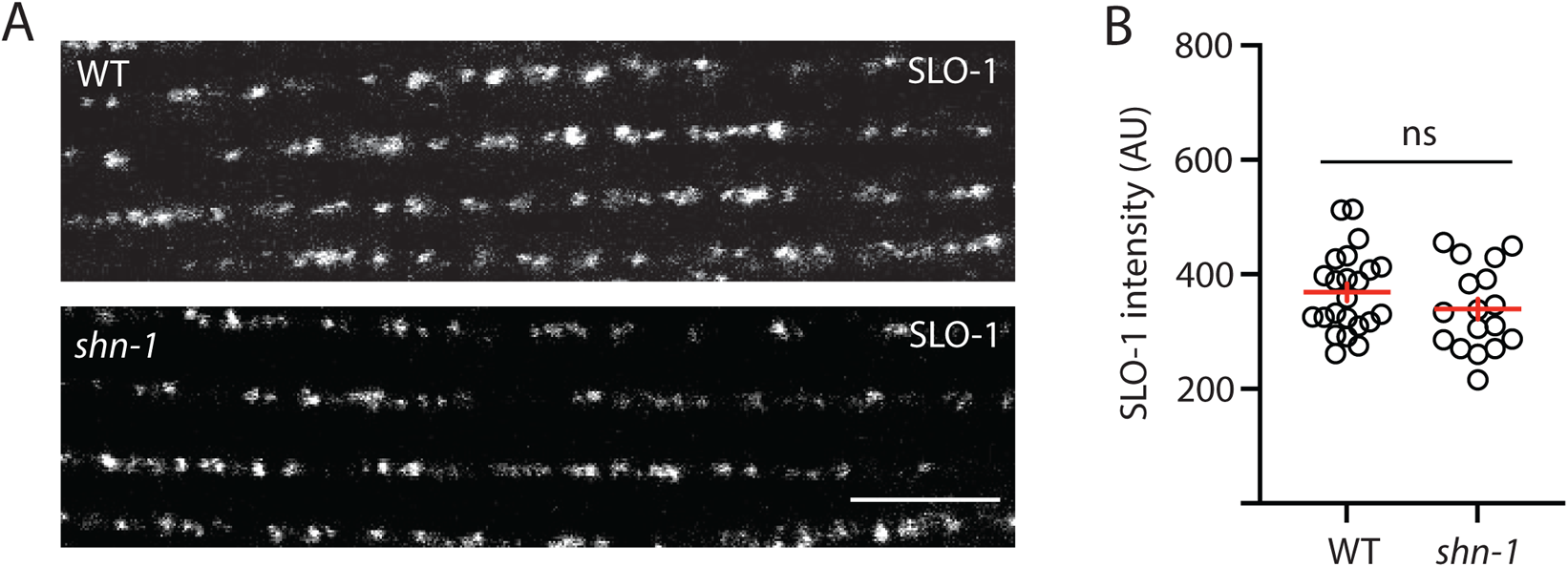
SLO-1 puncta intensity is unaltered in *shn-1* mutant muscles. SLO-1(nu678 GFP11) puncta intensity in body muscles is compared in WT and *shn-1(nu712* null) mutants. Representative images (A) and mean SLO-2 puncta intensity (B) are shown. Values that differ significantly from wild type controls are indicated (ns, not significant; *, *p* <0.05; **, *p* <0.01; ***, *p* <0.001). Error bars indicate SEM.

